# Resolution and b value dependent Structural Connectome in ex vivo Mouse Brain

**DOI:** 10.1101/2022.01.05.474963

**Authors:** Stephanie Crater, Surendra Maharjan, Yi Qi, Qi Zhao, Gary Cofer, James J. Cook, G. Allan Johnson, Nian Wang

## Abstract

Diffusion magnetic resonance imaging has been widely used in both clinical and preclinical studies to characterize tissue microstructure and structural connectivity. The diffusion MRI protocol for the Human Connectome Project (HCP) has been developed and optimized to obtain high-quality, high-resolution diffusion MRI (dMRI) datasets. However, such efforts have not been fully explored in preclinical studies, especially for rodents. In this study, high quality dMRI datasets of mouse brains were acquired at 9.4T system from two vendors. In particular, we acquired a high-spatial resolution dMRI dataset (25 μm isotropic with 126 diffusion encoding directions), which we believe to be the highest spatial resolution yet obtained; and a high-angular resolution dMRI dataset (50 μm isotropic with 384 diffusion encoding directions), which we believe to be the highest angular resolution compared to the dMRI datasets at the microscopic resolution. We systematically investigated the effects of three important parameters that affect the final outcome of the connectome: b value (1000 s/mm^2^ to 8000 s/mm^2^), angular resolution (10 to 126), and spatial resolution (25 µm to 200 µm). The stability of tractography and connectome increase with the angular resolution, where more than 50 angles are necessary to achieve consistent results. The connectome and quantitative parameters derived from graph theory exhibit a linear relationship to the b value (R^2^ > 0.99); a single-shell acquisition with b value of 3000 s/mm^2^ shows comparable results to the multi-shell high angular resolution dataset. The dice coefficient decreases and both false positive rate and false negative rate gradually increase with coarser spatial resolution. Our study provides guidelines and foundations for exploration of tradeoffs among acquisition parameters for the structural connectome in ex vivo mouse brain.

## Introduction

Diffusion magnetic resonance imaging (dMRI) is an important and unique technique to study brain microstructure and connections due to its sensitivity to random thermal motions of water (Alexander et al., 2010; Basser et al., 1994; Basser et al., 2000; Vu et al., 2015; Zhang et al., 2012). Diffusion tensor imaging (DTI), based on a Gaussian water displacement distribution, is the most commonly used approach because of its simplicity and its ability to afford insights to tissue characteristics (Basser et al., 1994). DTI only reconstructs a single direction of diffusion in each voxel, and inaccurately estimates fiber orientations in areas that contain multiple fiber populations (Frank, 2002; Le Bihan et al., 2001). A number of model-based and model-free methods have been developed to estimate the orientations of multiple axon populations per voxel, requiring a large number of diffusion-weighted images (DWIs) at the same b value (single-shell), or different b values (multi-shell) (Anderson, 2005; Assaf et al., 2008; Behrens et al., 2007; Hagmann et al., 2006a; Hutchinson et al., 2017; Tournier et al., 2004; Yeh et al., 2010b; Zhang et al., 2011). This acquisition method, termed as High Angular Resolution Diffusion Imaging (HARDI), has opened up new potentials to noninvasively measure microstructural features and study white matter connectivity (Berman et al., 2013; Descoteaux and Deriche, 2009; Frank, 2001; Le Bihan et al., 2006; Thomas et al., 2014; Tuch et al., 2002; Xie et al., 2015). Both tissue microstructure and connectome can be derived from the same dMRI dataset, whether the acquisition parameters optimized for tissue microstructure are still suitable for tractography are still unknown.

Due to the rapidly growing popularity of the HARDI technique for tractography, the influences of b value (strength of the diffusion gradient), angular resolution (number of gradient directions), and spatial resolution associated with tractography have been investigated in both human and monkey brains (Aydogan et al., 2018; Daianu et al., 2015; Jones et al., 2020; Schilling et al., 2017a; Schilling et al., 2018; Schilling et al., 2017b; Tournier et al., 2013). White matter fiber orientations of monkey brain were sensitive to both the number of diffusion directions (angular resolution) and b value (Schilling et al., 2017b). It has been reported that there was negligible improvement in accuracy when the b value of the single shell was increased from 6,000 to 12,000 s/mm^2^, or when the number of directions was increased from 64 to 96 (Schilling et al., 2018). Jones et al. reported that single-shell dMRI data can yield the same accuracy as multi-shell data for axonal orientation measurements if the reconstruction approach was chosen carefully (Jones et al., 2020). The accuracy remained stable across dMRI voxel sizes of 1 mm or smaller but degraded at 2 mm, particularly in areas of complex white-matter architecture (Jones et al., 2020). These studies provide quantitative measures of the reliability and limitations of dMRI acquisition and reconstruction methods, and can be used to identify the advantages of competing approaches as well as potential strategies for improving accuracy for fiber orientation distributions (Li et al., 2016; Maier-Hein et al., 2017). However, most of these studies focus on small areas of the white matter. Alterations due to the different acquisition parameters to the whole brain structural connectome are not fully investigated (Anderson et al., 2020; Caiazzo et al., 2018).

High spatial resolution dMRI has been developed to validate advanced fiber orientation distribution reconstruction methods with light microscopy imaging methods (Lefebvre et al., 2017; Mollink et al., 2017; Roebroeck et al., 2019). The high spatial resolution dMRI of human specimens has been achieved at 250 μm isotropic resolution, where each voxel still contains thousands of neurons (Jones et al., 2020). In contrast, the spatial resolution can be improved for the rodent brain in small-bore preclinical magnets with high-performance gradient coil, close-fitting radiofrequency (RF) coils, and optimized MR pulse sequences (Calabrese et al., 2015; Roebroeck et al., 2019; Wang et al., 2018a; Wu and Zhang, 2016). The dMRI of ex vivo postnatal mouse brains provided excellent contrasts revealing the evolutions of mouse forebrain structures at the spatial resolution of 94 x 96 x 93 μm^3^ (Zhang et al., 2005). With the isotropic spatial resolution of 52 μm, Aggarwal et al. were able to visualize the complex microstructure of embryonic cerebral tissue and resolve its regional and temporal evolution during cortical formation (Aggarwal et al., 2015). High-resolution diffusion MRI and tractography can reveal macroscopic neuronal projections in the mouse hippocampus and are important for future development of advanced tractography methods (Wang et al., 2020; Wu and Zhang, 2016).

In this study, we acquired HARDI datasets in ex vivo mouse brains on different MRI platforms (Varian platform and Bruker platform). In particular, we acquired a high-spatial resolution dMRI dataset (25 μm isotropic with 126 DWIs), which we believe to be the highest spatial resolution yet obtained (Wang et al., 2020); and a high-angular resolution dMRI dataset (50 μm isotropic with 384 DWIs), which we believe to be the highest angular resolution compared to the dMRI datasets at the microscopic resolution (Alomair et al., 2015; Aydogan et al., 2018; Johnson et al., 2019; Wu et al., 2013; Wu and Zhang, 2016; Yon et al., 2020). We systematically investigated the effects of three important parameters to the final connectome outcome: spatial resolution, angular resolution, and b value, in order to provide a foundation for exploring the tradeoffs among these acquisition parameters for the structural connectome in rodents.

## Methods

### Animal Preparation

Animal experiments were carried out in compliance with the Duke University Institutional Animal Care and Use Committee. Four wild-type adult C57BL/6 mice (Jackson Laboratory, Bar Harbor, ME) were chosen for MR imaging: 3 mice were scanned using Varian scanner and 1 mouse was scanned using Bruker scanner. For Varian scanner, one mouse was scanned at 25 μm isotropic resolution with b value of 3000 s/mm^2^; one mouse was scanned with 8 shells at 50 μm isotropic resolution (Wang et al., 2019); one mouse was scanned with 3 shells at 50 um isotropic resolution. For Bruker scanner, one mouse was scanned with 2 shells at 100 um isotropic resolution. Animals were sacrificed and perfusion fixed with a 1:10 mixture of ProHance-buffered (Bracco Diagnostics, Princeton, NJ) formalin. Specimens were immersed in buffered formalin for 24 hours and then moved to a 1:200 solution of ProHance/saline to shorten T1 (to about 110 ms) and reduce scan time (Wang et al., 2018a).

### MR histology protocol

MR images of 3 mice were acquired on a 9.4T Oxford 8.9-cm vertical bore magnet (Oxford Instruments, Abingdon, United Kingdom) with an Agilent VnmrJ 4.0 imaging console. A homemade solenoid RF coil was used for the scans. We used a three-dimensional (3D) diffusion-weighted spin-echo pulse sequence that was modified to support k-space undersampling and varying b values with different gradient directions. A 3D compressed sensing (CS) strategy fully sampled the readout dimension and under sampled the phase encoding dimensions using a sparsifying approach, which has been described in detail previously (Wang et al., 2018a). Acceleration factors (AF) of 1.0, 5.12 and 8.0 were used in this study, where 1.0 stands for the fully sampled data. To achieve multi-shell diffusion MRI acquisition, the diffusion gradient orientations were determined using the method proposed by Koay et al. to ensure the uniformity within each shell while maintaining uniformity across all shells (Koay et al., 2012).

Four protocols were developed in this study, three were on Varian platform and one was on Bruker platform. The three protocols used for Varian platform were: 1) An under sampled dataset (AF = 8.0) was acquired using a single-shell acquisition (b value = 3000 s/mm^2^) at 25 μm isotropic resolution, 126 DWIs and 13 b0 images, and one repetition. Total scan time was about 127 hours. 2) An under sampled dataset (AF = 5.12) was acquired with 8 shells (b value = 1000, 2000, 3000, 40000, 5000, 6000, 70000, 8000 s/mm^2^) at 50 μm isotropic resolution, 48 DWIs and 4 b0 images in each shell, and one repetition. The total scan time was 148 hours. 3) An under sampled dataset (AF = 5.12) was acquired with 3 shells (1500, 4000, and 8000 s/mm^2^) at 50 μm isotropic resolution, 61 DWIs and 6 b0 images in each shell, and one repetition. The total scan time was 71.5 hours. Temperature was monitored throughout all the scans and fluctuation was less than 1 °C. All the scans kept the same TE of 15.2 ms and TR of 100 ms. Gradient separation time was 7.7 ms, the diffusion gradient duration time was 4.8 ms, and the maximum gradient amplitude was 134 G/cm.

MR images of one mouse brain was acquired on a 30-cm bore 9.4T magnet (Bruker BioSpec 94/30, Billerica, MA). A high-sensitivity cryogenic RF surface receive-only coil was used for signal reception (Bruker CryoProbe). The protocol used for Bruker platform was: a three-dimensional (3D) diffusion-weighted spin-echo pulse sequence was used for imaging acquisition with the scan parameters: two shells (b value = 3,000 and 8,000 s/mm^2^), 100 μm isotropic resolution, 61 DWIs and 6 b0 images in each shell, and one repetition. The total scan time was 41 hours. Details of the acquisition parameters are summarized in Table 1.

**Table 1:** The acquisition parameters of 4 different diffusion MRI datasets.

| Resolution<br>($\mu\text{m}$) | Shells | b-value<br>($\text{s/mm}^2$) | Gradient<br>Orientations | TR<br>(ms) | TE<br>(ms) | FOV | Matrix Size | Scan<br>time | Scan<br>time/<br>volume | Mice<br>Numbers | Scanner |
| --- | --- | --- | --- | --- | --- | --- | --- | --- | --- | --- | --- |
| 50 | 8 | 1k,2k,3k,4k,<br>5k,6k,7k,8k | 48/shell | 100 | 15.2 | 19.2*12.8*12.8 | 384*256*256 | 148 h | 0.35 h | 1 | Varian |
| 25 | 1 | 3k | 126/shell | 100 | 15.2 | 19.2*12.8*12.8 | 768*512*512 | 127 h | 0.91 h | 1 | Varian |
| 50 | 3 | 1.5k, 4k, 8k | 61/shell | 100 | 15.2 | 19.2*12.8*12.8 | 384*256*256 | 71.5 h | 0.36 h | 1 | Varian |
| 100 | 2 | 3k, 8k | 61/shell | 100 | 23.8 | 18.0*12.8*7.6 | 180*128*76 | 41 h | 0.31 h | 1 | Bruker |

To assess the effect of angular resolution, the diffusion dataset acquired using protocol 1 was subsampled from 10 to 110 gradient orientations. To assess the effect of spatial resolution, the same dataset was also down sampled to 50, 100, and 200 μm isotropic resolution in k-space. To assess the effect of b value, the diffusion dataset acquired using protocol 2 was subsampled to single shell (from 1000 to 8000 s/mm^2^), two shells (1000 and 2000 s/mm^2^, 3000 and 4000 s/mm^2^, 5000 and 6000 s/mm^2^, 7000 and 8000 s/mm^2^), and the various multi-shell combinations up to 7 shells. Reconstruction of the under sampled k-space data has been described in previous studies (Hollingsworth, 2015; Lustig et al., 2007; Wang et al., 2018b). In particular, acceleration factors of 5.12 and 8.0 indicate sampling of only 1/5.12^th^ and 1/8^th^ of the fully sampled data. For example, the scan time for protocol 1 took 127 hours; the scan time for the fully sampled dataset would take 127*8.0 = 1016 hours (more than 42 days). The reconstruction was implemented on a Dell Cluster where an initial Fourier transform along the readout axis yielded multiple 2D images from each 3D volume which could be spread across the cluster for parallel reconstruction. The image single-to-noise ratio (SNR) was calculated as the ratio of the average signal value to the standard deviation of the background.

### Diffusion metrics and Tractography

Firstly, all the DWIs were registered to the baseline image (b0). Diffusion tensor imaging (DTI) was then used to caluclate the tensor and the scalar indices: AD (axial diffusivity), RD (radial diffusivity), FA (fractional anisotropy), and MD (mean diffusivity). Tractography of generalized Q-sampling imaging (GQI) was obtained by the streamline tracking algorithm implemented in DSI Studio with maximum four fibers resolved in one voxel (Yeh et al., 2010a; Yeh et al., 2013). The multiple fiber ratio (MFR) at each b value or spatial resolution is defined as: 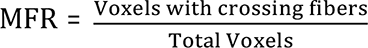, where the Total Voxels are the voxels of the whole mouse brain. The propagation process was repeated until the tracking trajectory exceeded either a turning angle of greater than 45°, or the anisotropy value of the current position was below a predefined threshold of normalized quantitative anisotropy (0.075). Ten million fibers were generated with whole brain seeding and the mean tract length of the whole brain tractography was calculated. Tract density imaging (TDI) was also used to facilitate visualization of the whole brain microstructures after tractography is generated (Calamante et al., 2012; Yeh et al., 2013). All fiber tracking operations were preformed using the DSI studio toolbox (Yeh et al., 2013). To quantitatively evaluate the performance of tractography and connectome results, the network topology properties were measured using Brain Connectivity Toolbox (Rubinov and Sporns, 2010). Furthermore, three quantitative parameters are also calculated: false positive rate 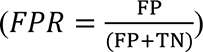, false negative rate 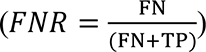, and dice coefficient 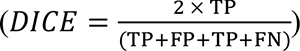, where TP is true positives, FP is false positives, TN is true negatives, and FN is false negatives (Aydogan et al., 2018). The threshold of the connectivity map for DICE ranged from 0.001 to 0.2; the threshold of the FPR and FNR was 0.001. The full dMRI dataset from protocol 1 (25 μm with 126 DWIs) was used as ground truth (GT) for assessing the effect of spatial resolution and angular resolution; The full dMRI dataset from protocol 2 (50 μm with 384 DWIs) served as ground truth for assessing the b value effect. The diffusion MRI metrics are available from: https://civmvoxport.vm.duke.edu upon request. To validate the fiber orientation distribution at each voxel for the brain, the diffusion MRI dataset with 3 different b values (1500, 4000, and 8000 s/mm^2^) was also analyzed using the constrained spherical deconvolution (CSD) method provided by MRTrix3 (https://www.mrtrix.org/) with a maximum harmonic order of 6. The fiber orientation distribution at six regions of the brain were compared at different b values.

### Comparison with Allen Mouse Brain Atlas Neuronal Tracer data

The detailed tracer injection and projection density images derived from the Allen Mouse Brain Atlas (AMBA) can be served as the gold standard to validate the diffusion tractography results (Oh et al., 2014). The tracer injection and projection density images were downloaded from AMBA (25 µm resolution) (https://connectivity.brain-map.org/) and converted to NIFTI format. Five injections were selected from different parts of the brain with the injection IDs of 113166056, 127139568, 174957972, 307297141, and 309385637. The diffusion MRI images at 25 µm isotropic resolution with 126 DWIs were registered to the AMBA using Advanced Normalization Tools (ANTs) automated image registration, which has been described in previous study in detail (Avants et al. 2011; Calabrese et al., 2015). The 5 injections sites were then transformed back to MRI space as the seeding regions for tractography and TDI images. The projection density images were also transformed back to MRI space and directly compared with the TDI images using dice coefficient measurement.

## Results

Figure 1 shows representative DTI metrics and DWIs at different b values (0 – 8000 s/mm^2^). In Figure 1a, the corpus callosum exhibits a higher FA value compared to cortex region, while the MD value shows the opposite trend. The signal intensity gradually decreased with higher b value, the image at b value of 8000 s/mm^2^ still exhibits relatively high SNR (Fig1b-1c). The image magnitudes are normalized to the b0 image intensity. The SNR values (Fig1d) varies from 85 (without diffusion weighting) to 15 (b value of 8000 s/mm^2^).

**Figure 1:**
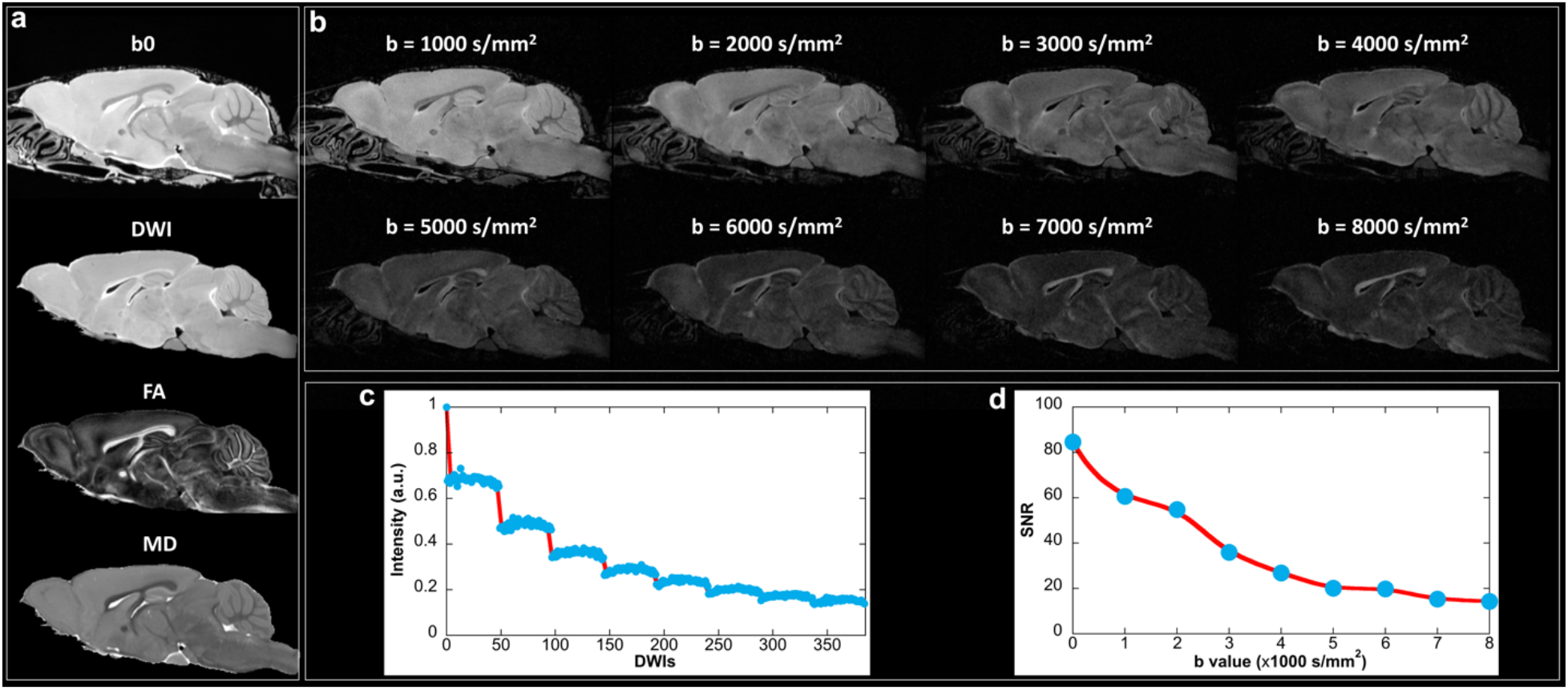
Representative DTI metrics (a) and DWIs (b) at different b values (0 – 8000 s/mm^2^). The white matter exhibited a higher FA value compared to gray matter, while the MD value showed the opposite trend. The signal intensity gradually decreased with higher b value, the image at b value of 8000 s/mm^2^ still exhibits relatively good image quality. The image magnitudes are normalized to the b0 image intensity. The SNR values varies from 85 (without diffusion weighting) to 15 (b value of 8000 s/mm^2^).

Figure 2 shows the structural connectivity maps at different threshold (0.0001 – 0.2), where connections become sparser with higher threshold (Fig2a-2d). The dice coefficient is strongly dependent on b values for both single-shell (Fig2e) and multi-shell acquisitions (Fig2f-2h). The highest dice coefficient is found at b value of 3000 s/mm^2^ with single-shell acquisition regardless of the threshold. High dice coefficient is found at b value of 3000 and 4000 s/mm^2^ as well as 5000 and 6000 s/mm^2^ with two-shell acquisition. The dice coefficient is consistently high (> 0.85) using more than two shells (Fig2g-2h). The false negative ratio (FNR) gradually decreases with higher b value, while the false positive ratio (FPR) gradually increases with b value (Fig2i). The crossing point of these two curves occurs at the b value of about 3000 s/mm^2^. The similar trend has been found for two-shell acquisitions (Fig2j). Both FPR and FNR are lower than 0.2 when more than two shells are applied (Fig2k-2l). In general, higher dice coefficient, lower FPR, and lower FNR are found with more shells added (Supplemental Figure 1).

**Figure 2:**
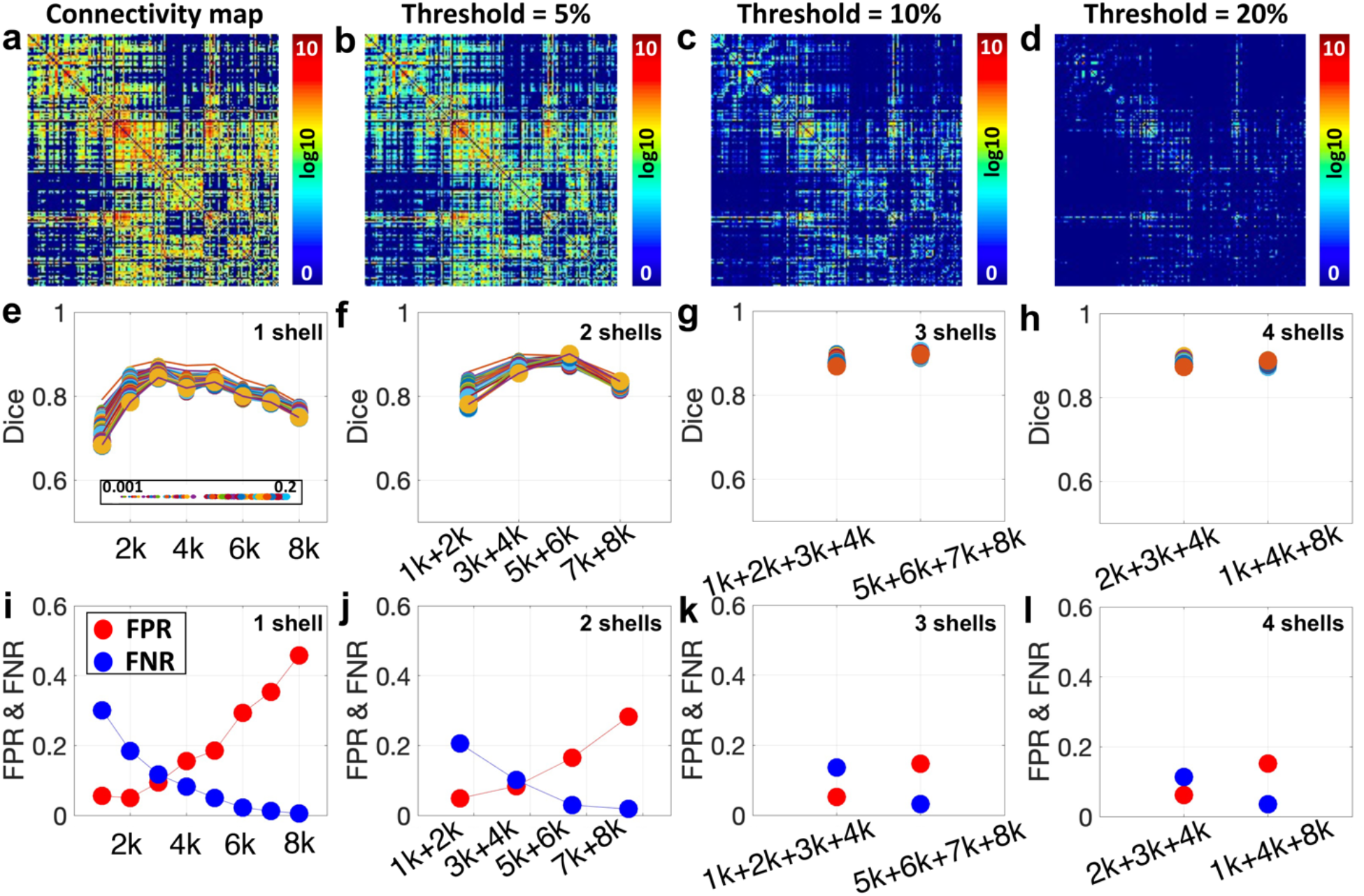
The connectivity maps at different threshold (0.0001 – 0.2), where connections become sparser with higher threshold (a-d). The highest dice coefficient is found at b value of 3000 s/mm^2^ with single-shell acquisition (e) regardless of the threshold (0.0001 – 0.2). The false negative ratio (FNR) gradually decreases with higher b value, while false positive ratio (FPR) gradually increases with b value (i, j). Both FPR and FNR are lower than 0.2 using with more than two shells (k, l).

The FPR and FNR variations with b value are also demonstrated in individual tracts (Figure 3), including hippocampus (Hc), primary motor cortex (M1), and corpus callosum (cc). More FN tracts are apparent at lower b value (green arrows), while more FP tracts are seen at higher b values (white arrows). The tracts at b value of 3000 s/mm^2^ are visually comparable to the ground truth, although a few FP and FN tracts can still be observed. The tracts from multi-shell (2000, 3000, and 4000 s/mm^2^) acquisition show good agreement with GT (purple arrows). The complex fiber orientation distributions at different brain areas are demonstrated in Supplemental Figure 2. The consistent results are also evident in the structural connectome of isocortex and white matter (Figure 4). Compared to the GT, there are more FP connections at higher b value (8000 s/mm^2^), while the connections become sparser at b value of 1000 s/mm^2^.

**Figure 3:**
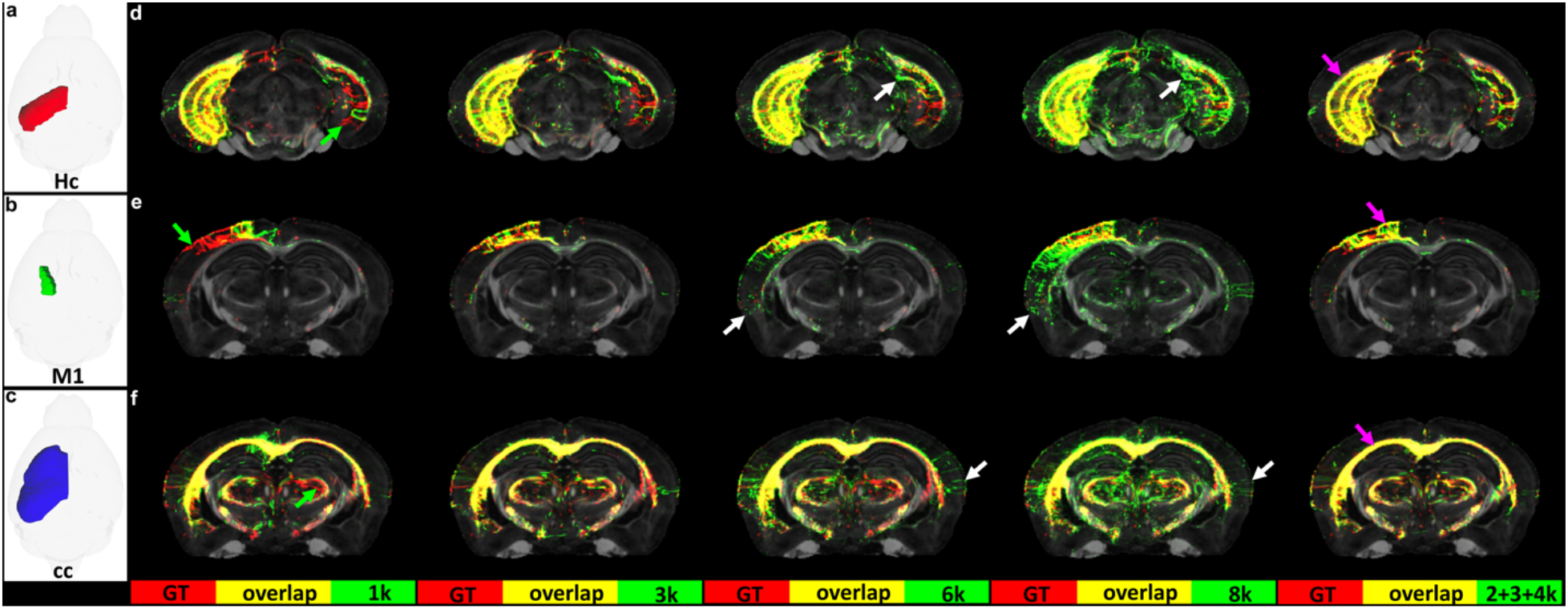
The tract density imaging (TDI) at different b values, including hippocampus (Hc), primary motor cortex (M1), and corpus callosum (cc). Compared to the ground truth (TDI in red color), the false negative (FN) tracts are apparent at lower b value (green arrows), while more false positive (FP) tracts exist at higher b values (white arrows). The tracts at b value of 3000 s/mm^2^ are visually comparable to the ground truth, although a few false positive (FP) and false negative (FN) tracts can still be observed. The tracts from multi-shell (2000, 3000, and 4000 s/mm^2^) acquisition also show good overlap with the ground truth (purple arrows).

**Figure 4:**
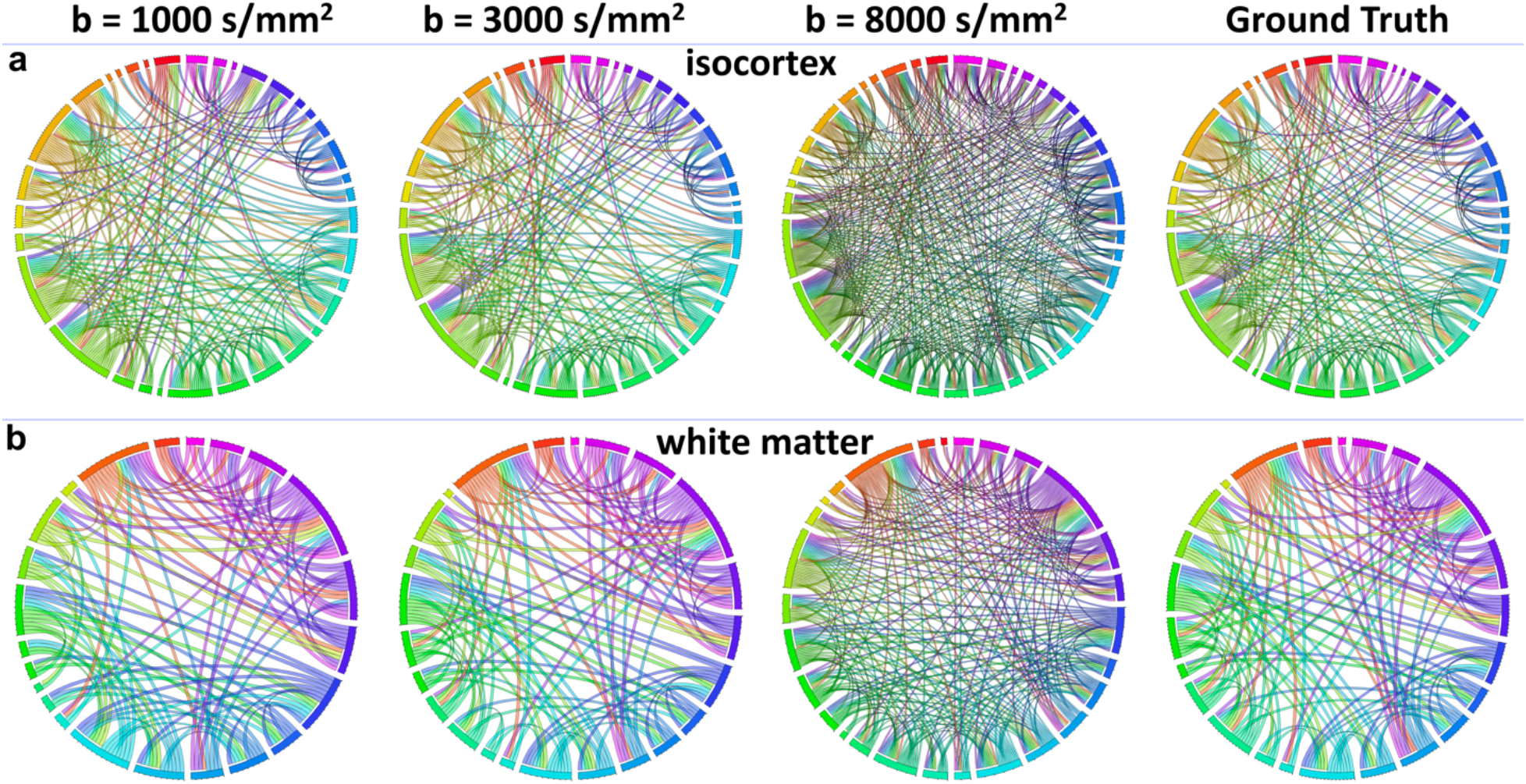
Compared to the ground truth (GT), there are more false positive (FP) connections in isocortex and white matter at higher b value (8000 s/mm^2^), while the connectome becomes sparser at b value of 1000 s/mm^2^.

In order to explore the b value dependent connectome, the MFR mapping is illustrated in Figure 5, where the GT is from the dMRI dataset with all the b values (1000-8000 s/mm^2^, 384 DWIs). The cortex is populated with multiple crossing fibers and the cc and fimbria (fi) are dominated by single fiber, while the olfactory bulb (Ob), Hc, and cerebellum contain both crossing fibers and single fiber. The MFR is extremely low at b value of 1000 s/mm^2^, which suggests that crossing fibers cannot be well resolved at this b value. The MFR gradually increases with b value and the MFR is close to 1 (1 means every voxel of the brain contains crossing fibers) at b value of 8000 s/mm^2^ (Fig5q). For the single-shell acquisition, the MFR at b value of 3000 s/mm^2^ shows the highest agreement with GT. The MFR becomes more stable when more shells are added (Fig5r-5t). The b value dependent fiber orientation distributions in Hc and cc regions are illustrated in Supplemental Figure 3 and Supplemental Figure 4.

**Figure 5:**
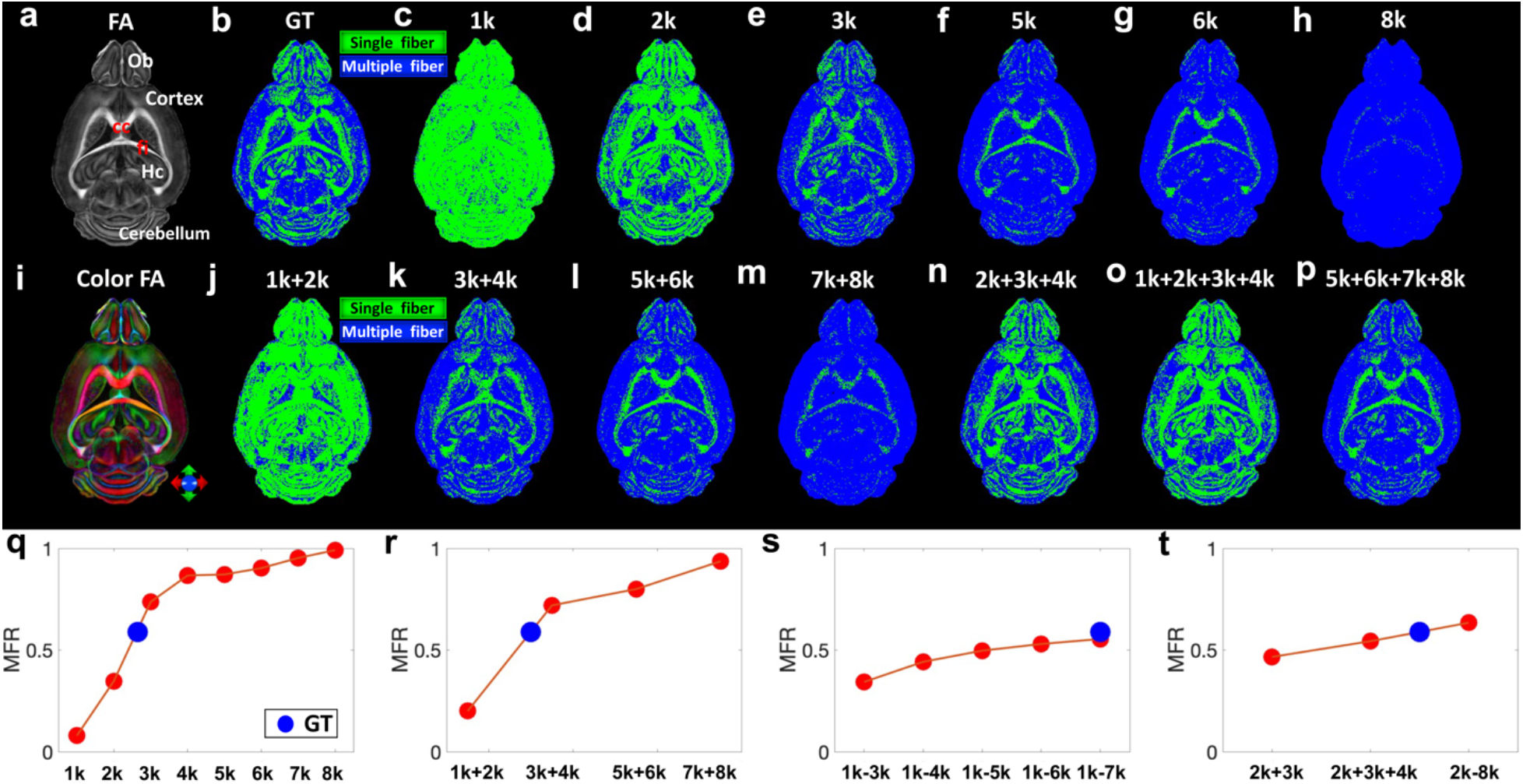
Multiple fiber ratio (MFR) mapping at different b values. The ground truth (GT) is the dataset with all the b values (1000 - 8000 s/mm^2^, 384 DWIs). The MFR is extremely low at b value of 1000 s/mm^2^. The MFR gradually increases with b value and the MFR is close to 1 (1 means every voxel of the brain has crossing fibers) at b value of 8000 s/mm^2^ (h, q). For the single-shell acquisition, the MFR at b value of 3000 s/mm^2^ matches with the ground truth best (q). The MFR is more stable when more shells are added (r, s, t). Green color: single fiber; Blue color: Multiple fibers; Ob: olfactory bulb; cc: corpus callosum; fi: fimbria; Hc: hippocampus.

The MFR mapping of a multi-shell (b value of 3000 s/mm^2^ and 8000 s/mm^2^) dMRI dataset acquired on Bruker platform is shown in Figure 6. Compared to the fiber orientation distributions at lower b value (3000 s/mm^2^), the cc and Hc are dominated by the crossing fibers at b value of 8000 s/mm^2^ (black and white arrows in Fig6b-6c). Few voxels through the whole brain contain single fiber at higher b value, while more voxels exhibit single fiber at lower b value (Fig6d-6f). The similar findings are demonstrated with the multi-shell dMRI dataset (b value of 1500, 4000, and 8000 s/mm^2^) using both MRtrix3 and DSI studio (Supplemental Figure 5 and Supplemental Figure 6).

**Figure 6:**
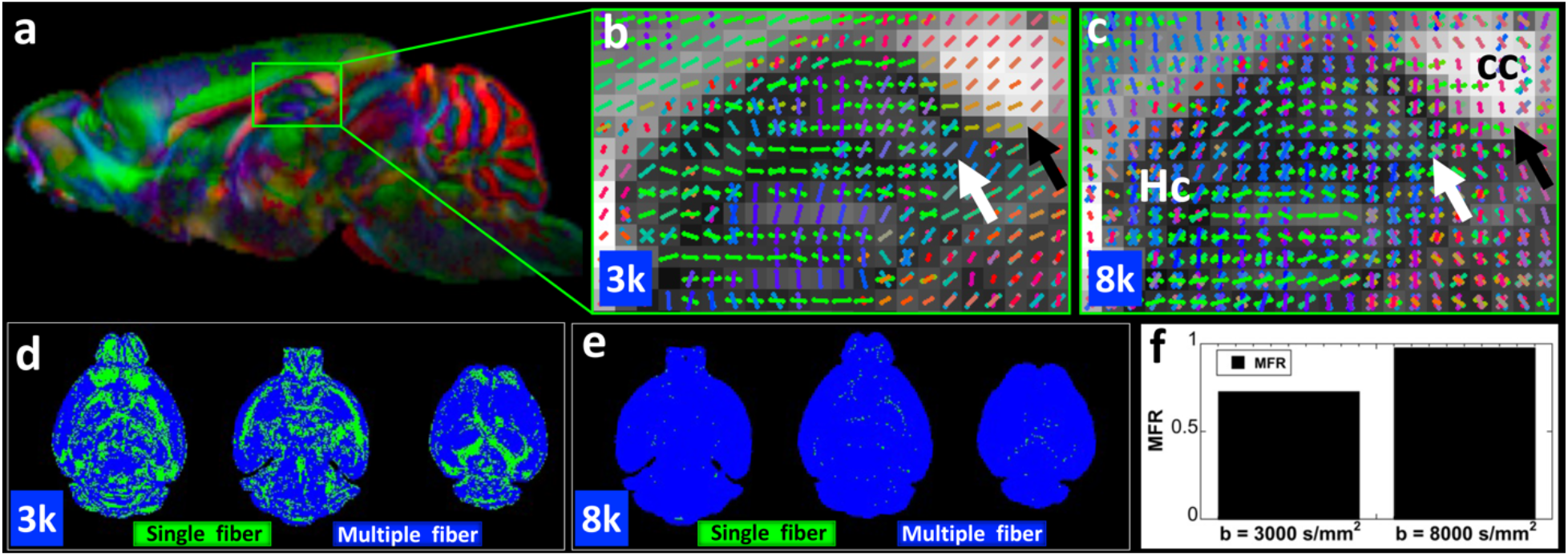
Multiple fiber ratio (MFR) mapping at different b values (3000 and 8000 s/mm^2^) acquired using Bruker scanner. The cc and Hc (b, c) are dominated by crossing fibers at higher b value (8000 s/mm^2^). The MFR of the whole brain is close to 1 (f) at the b value of 8000 s/mm^2^. cc: corpus callosum; Hc: hippocampus.

To quantitatively compare the b value dependent structural connectome, the metrics from graph theory has been plotted as the function of the b value (Figure 7). Linear relationship between network parameters and b values has been observed for all the three parameters (R^2^ > 0.99), including density, path length, and small-world. For single-shell acquisition, the quantitative values at b value of 3000 s/mm^2^ matches well with the GT (Fig7a-7c, purple dots). For two-shell acquisition, the quantitative values at b value of 3000 and 4000 s/mm^2^ show consistent results with the GT (Fig7d-7f).

**Figure 7:**
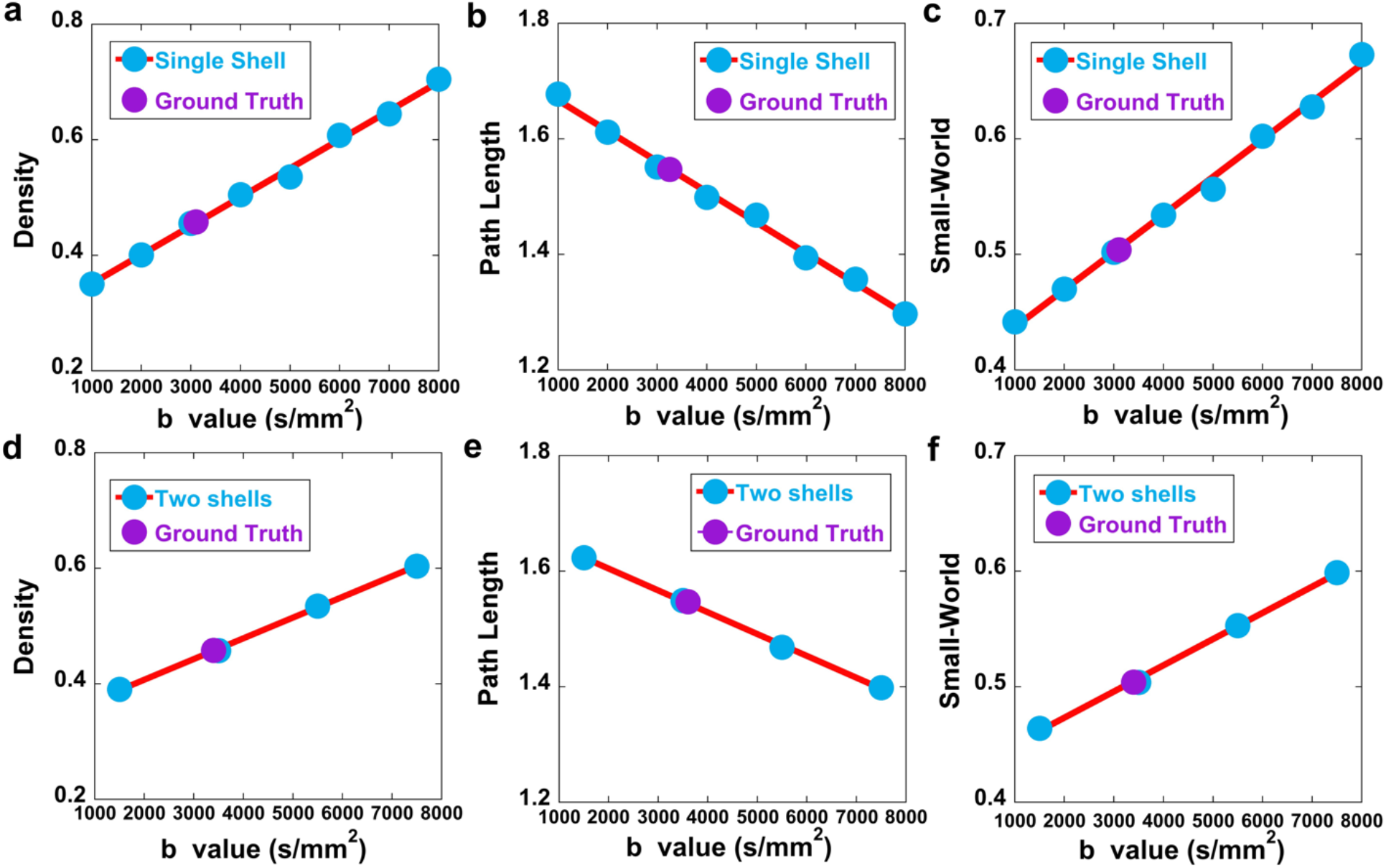
The quantitative metrics from graph theory as a function of the b value. Linear relationship between Graph Theory and b value has been observed for all three parameters (R^2^ > 0.99), including density, path length, and small-world. For single-shell acquisition, the quantitative values at b value of 3000 s/mm^2^ matches well with the GT (a-c, purple dots). For two-shell (1000 and 2000 s/mm^2^, 3000 and 4000 s/mm^2^, 5000 and 6000 s/mm^2^, 7000 and 8000 s/mm^2^) acquisition, the quantitative values at b value of 3000 and 4000 s/mm^2^ show consistent results with the GT (d- f).

The brain structural connectome is also strongly dependent on the angular resolution, which is demonstrated in Figure 8. The whole brain connectome become sparser at lower angular resolution (Fig8a-8f). Two distinct growth stages of the dice coefficient curves are shown as a function of the angular resolution (Fig8g). A steep increase stage when the angular resolution is lower than 50 (0.4 < dice coefficient < 0.85), and a stable stage when the angular resolution is higher than 50 (dice coefficient > 0.85). The FNR drops dramatically when the angular resolution increases from 10 (∼ 0.6) to 50 (∼ 0.15) and becomes more stable when the angular resolution is higher than 50 (Fig8h). In contrast, the FPR drops dramatically with more angles and becomes more stable when the angular resolution is higher than 50 (Fig8h). The variation in graph theory parameters (small-world, global efficiency, and clustering coefficient, Fig8i) respect to the angular resolution is similar to the dice coefficient (Fig8g), with an increase in all three as the angular sampling increases from 10 to 50 and less change for angular sampling above 50.

**Figure 8:**
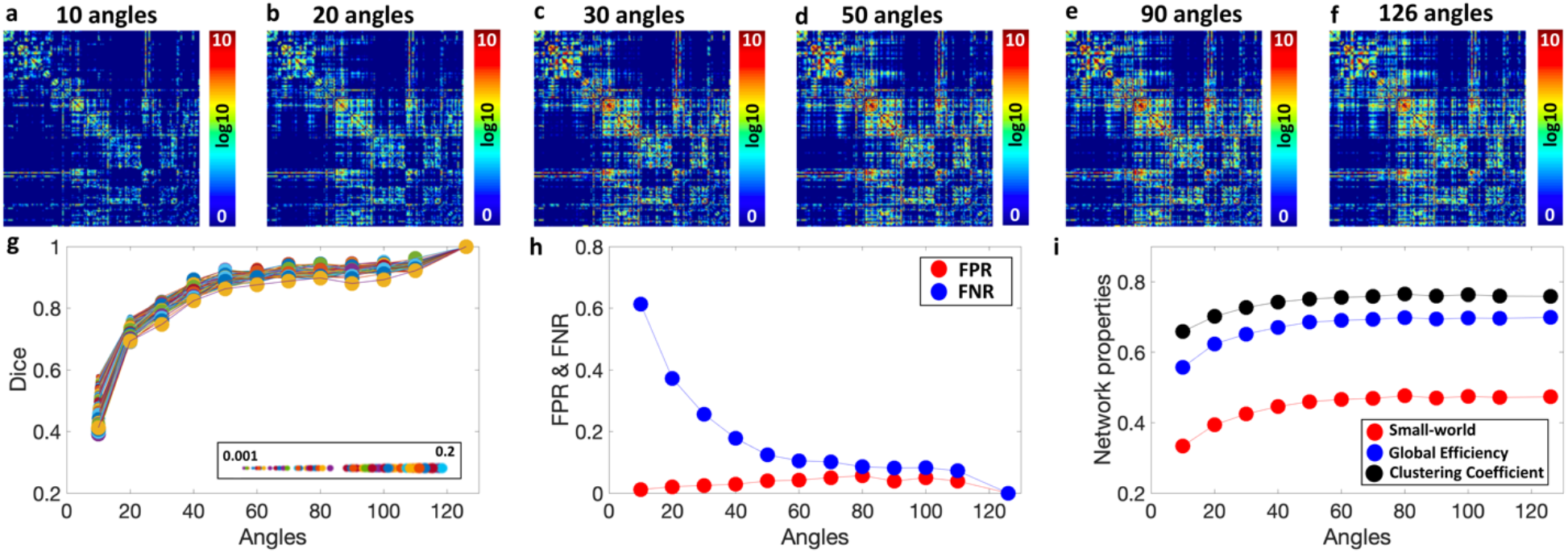
The angular-dependent brain structural connectome. The whole brain connectome becomes sparser at lower angular resolution (a-f). The false negative ratio (FNR) drops dramatically with more angles and becomes more stable when the angular resolution is higher than 50 (h). In contrast, the false positive ratio (FPR) remains at a low level (< 0.1) through all the angular resolutions. The quantitative parameters (small-world, global efficiency, and clustering coefficient, i) variations respect to the angular resolution is similar to the dice coefficient (g), where more dramatically change is seen when the angular resolution is lower than 50.

Figure 9 shows the tractography results of right corpus callosum (Fig9a-9d) and right hippocampus (Fig9e-9h) at different angular resolution (10, 60, and 126 DWIs). The interhemispheric connections (red and green arrows) and long-range connections (white arrows) are largely reduced at lower angular resolution (Fig9b, 9f). The tract length graduate increases with angular resolution and becomes more stable when the angular resolution is higher than 50 (Supplemental Figure 7).

**Figure 9:**
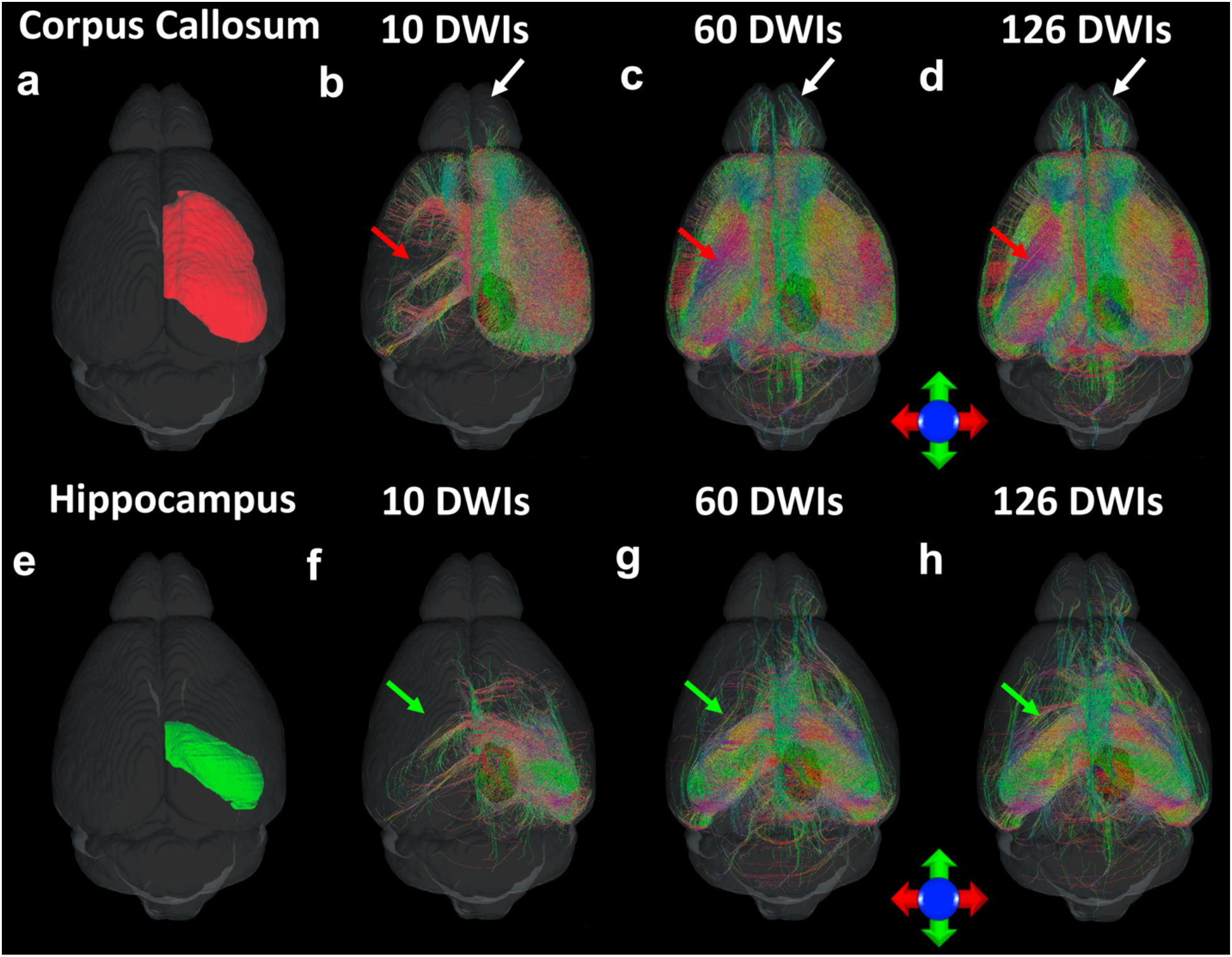
The tractography result of the right corpus callosum (a-d) and right hemisphere (e-h) at different angular resolution (DWIs of 10, 60, and 126). The interhemispheric connections (red and green arrows) and long-range connections (white arrows) are largely reduced at lower angular resolution. DWI: diffusion weighted imaging.

Figure 10 shows the impact of the spatial resolution on the structural connectome. The neuroanatomy of the cerebellum is highlighted in the FA images at spatial resolution ranging from 25 µm to 200 µm. The microstructure and fiber directions are better resolved in white matter (WM), molecular layer (ml), and granular layer (gl) at higher spatial resolution (Fig10a). In contrast, these structures cannot be well resolved at lower spatial resolution (Fig10d). The dice coefficient decreases with lower spatial resolution, while both FPR and FNR gradually increase with lower spatial resolution. The capability of resolving crossing fibers is also reduced at lower spatial resolution, where MFR drops from ∼ 0.8 (25 µm) to ∼ 0.6 (100 and 200 µm).

**Figure 10:**
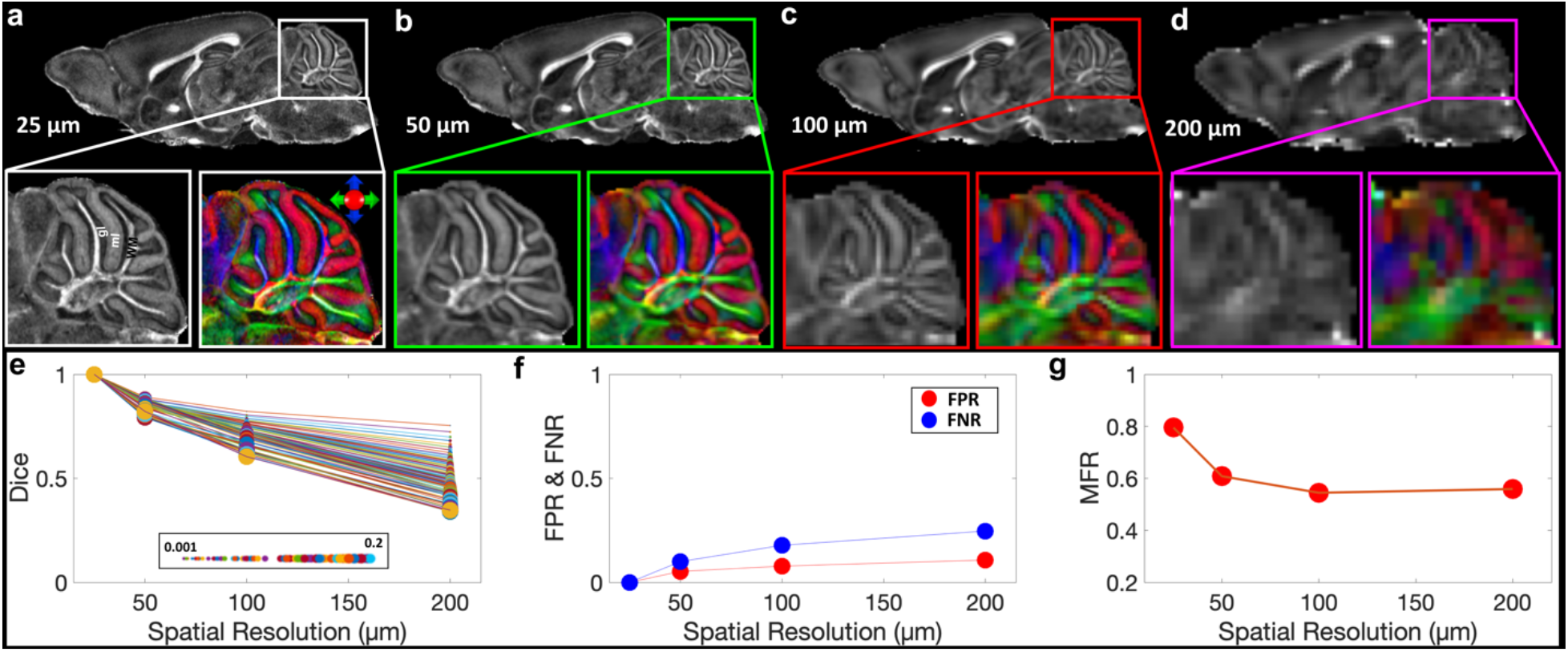
The effect of the spatial resolution to the brain connectome. The microstructure and fiber directions are better resolved in white matter (WM), molecular layer (ml), and granular layer (gl) at higher spatial resolution (a-d). The Dice Coefficient decreases with lower spatial resolution, while both false positive ratio (FPR) and false negative ratio (FNR) gradually increase with lower spatial resolution. The capability of resolving crossing fibers is also reduced with lower spatial resolution.

The AMBA has been served as the gold standard to validate the diffusion tractography results. The overlap between tractography results from dMRI and projection density images from AMBA at the same spatial resolution (25 µm isotropic) are shown in Figure 11. The dice coefficient ranges from 03126 to 0.5871, depending on different injection sites. The inconsistencies between these two methods are illustrated in white and purple arrows.

**Figure 11:**
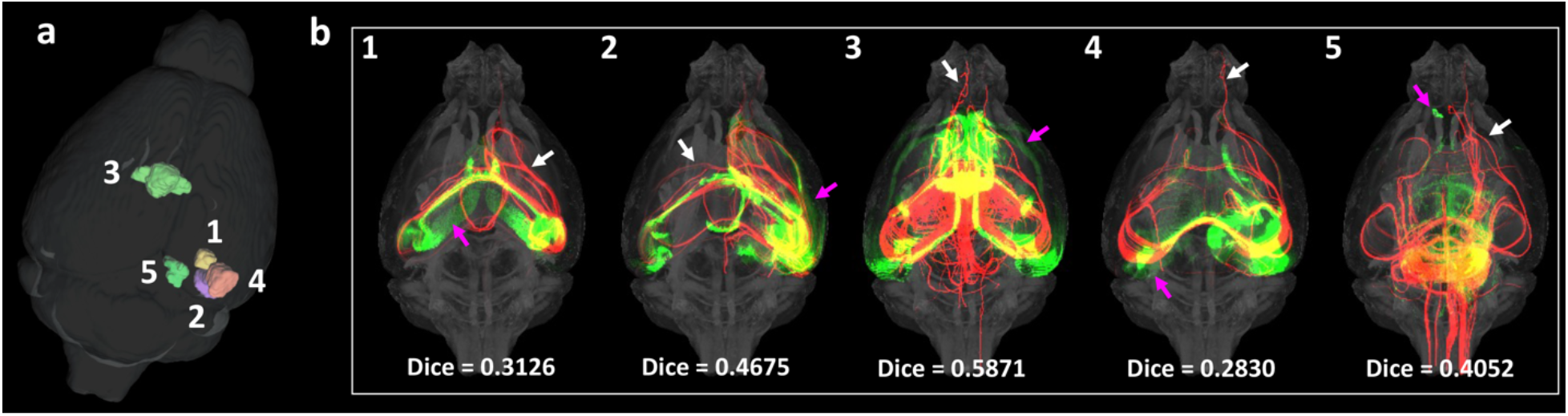
The overlap of the tracer projection density for 5 injection sites (green color) and tractography results (red color). The Dice coefficient varies from 0.2830 to 0.5871 at different injections sites.

## Discussion

The influences of experimental factors of dMRI associated with the tractography and connectome have been investigated in detail in primate brains, but not fully explored in rodents (Anderson et al., 2020; Caiazzo et al., 2018). Various biophysical models have been developed to probe the tissue microstructure, but the b values may vary dramatically for different models (Alexander et al., 2002; Assaf et al., 2008; Calamante et al., 2010; Edwards et al., 2017; Henriques et al., 2019; Huang et al., 2020; Kaden et al., 2016; Tournier et al., 2004; Tournier et al., 2011; Tuch et al., 2002; Wedeen et al., 2008; Zhang et al., 2011). The same dMRI dataset may be used for both microstructure and structural connectome analysis. However, whether the acquisition parameters optimized for tissue microstructure are still suitable for tractography are still unknown. The effects of different acquisition parameters to the final structural connectome are essential for setting up protocols for tractography and connectivity studies. To address these issues, we investigated the effects of three experimental parameters of dMRI scans: b value (1000 to 8000 s/mm^2^), spatial resolution (25 to 200 µm), and angular resolution (10 to 126 DWIs).

### The effect of b value

An important technical tradeoff for dMRI acquisition is performing adequate diffusion weighting to resolve crossing fibers while preserving sufficient SNR (Setsompop et al., 2013). The proper b value for a specific study is dependent on the tissue properties, diffusion models, and the gradient strength of MRI scanners (de Figueiredo et al., 2011; Hagmann et al., 2006b; Jelescu et al., 2015; Xie et al., 2015). The HCP scanner can achieve b values up to 20,550 s/mm^2^ with maximum gradients of 300 mT/m to generate structural connectome and probe the different tissue microstructures in live human (Duval et al., 2015; Guglielmetti et al., 2016; Jensen et al., 2005; Landman et al., 2007; Palombo et al., 2020; Zhang et al., 2012). The effect of b value on tractogram has been investigated in previous studies, but remains largely unanswered for preclinical MRI (Anderson et al., 2020; Schilling et al., 2017b; Sepehrband et al., 2017; Sotiropoulos and Zalesky, 2017). A relative lower b value may be adequate for accurately quantifying the DTI metrics, such as FA and MD. However, a lower b value is not sufficient for resolving crossing fibers in brain regions with complex tissue structures (Tournier et al., 2013). Higher b value is preferred to better resolve the crossing fibers, but also suffers from the lower SNR, which may fail to accurately resolve the FODs (Alexander and Barker, 2005; Yeh et al., 2010a). It has been reported that the depiction of crossing fiber by HARDI-based multi-tensor tractography was not substantially influenced by b values ranging from 700 to 2800 s/mm^2^. However, this study was conducted on a 1.5T MRI scanner and mainly focused on optic tracts (Akazawa et al., 2010). In a later study, Tournier et al. reported that the highest angular resolution achievable in practice is obtained using a b value of approximately 3000 s/mm^2^ to characterize the DW signal at 3T (Ferizi et al., 2017; Tournier et al., 2013).

In the preclinical domain, multi-shell data reconstructed major fibers with less error than single-shell data, and was most successful at reducing the angular error when the lowest shell was excluded in a mouse study (Daianu et al., 2015). We also demonstrated that the crossing fibers can barely be resolved at b value of 1000 s/mm^2^. In fact, a relative low b value is often used in dMRI studies to increase the SNR, distinguish different components of an imaging voxel, and fit different biophysical models (Ferizi et al., 2017; Hutchinson et al., 2017). For instance, FA and MD derived from the basic tensor model are known to be very sensitive to the b value (Tournier et al., 2011; Wu and Miller, 2017). A recent study showed that lower b value is essential to accurately estimate neurite orientation dispersion and density imaging (NODDI) metrics and significant artifacts were evident when fitting this biophysical model using only high b values (5000 – 8000 s/mm^2^) (Wang et al., 2019). In the current study, the MFR is higher than 0.9 at b value of 7000 s/mm^2^ (48 DWIs) or 8000 s/mm^2^ (48 DWIs). The MFR remains high (> 0.9) when these two shells are combined (96 DWIs). To accurately resolve the crossing fibers at higher b value, the angular resolution may need to increase dramatically to compromise the SNR reduction. In contrast, a lower b value (∼ 1000 s/mm^2^) which is crucial for probing tissue microstructure in various biophysical models may reduce the accuracy in estimating the fiber orientation distributions and generating structural connectome.

Although multi-shell HARDI acquisition has advantages for tractography and structural connectome, many studies are limited to single-shell acquisition due to limited scan time or the balance between spatial resolution and angular resolution. We demonstrated that the structural connectome is strongly dependent on the b value. A medium b value around 3000 s/mm^2^ may be a good option for resolving crossing fibers, generating tractography, and calculating connectivity maps, if only single-shell acquisition is conducted for the study. For the quantitative parameters derived from graph theory, it’s interesting to observe that these parameters exhibit linear relationship as a function of the b value. The dMRI protocols likely vary from different studies, vendors, and gradient sets, thus, quantitative comparison of structural connectome from multi-sites or multi-vendors is still challenging. The linear relationship between the network parameters and b value may help to mitigate this deviation even the datasets are acquired at different b values. However, this is only demonstrated for ex vivo mouse brain studies, whether it holds the similar promise need further exploration in both human and in vivo rodent studies (Caiazzo et al., 2018).

### The effect of angular resolution

High angular resolution becomes more important for dMRI because it is widely acknowledged that the basic tensor model is not appropriate in regions that contain multi fiber orientations (Hope et al., 2012; Jones, 2004). Many of these higher order models have been proposed, relying on the HARDI acquisition to achieve a relatively large number of DWIs images at the same b value or different b values. The number of directions required for tractography using DTI and higher order models have been discussed in previous studies in white matter tracts of human brain. It is recommended to acquire more than the minimum of 45 DW directions to avoid issues with imperfections in the uniformity of the DW gradient directions and to meet SNR requirements of the intended reconstruction method (Tournier et al., 2013). A more recently study, more than 45 DW directions were recommended to resolve the complex fibers and detect the number of fiber populations in a voxel more accurately in white matter area (Schilling et al., 2017b). In the mouse brain with 43 µm isotropic resolution, it has been also suggested that a dMRI protocol with more than 45 angular samples helped to substantially reduce the tractography errors (Anderson et al., 2020). In this study, both dice coefficient and network properties achieve plateaus when the angular resolution is higher than 50. Meanwhile, both FPR and FNR remains relatively low with more than 50 angles. Thus, 50 DWIs or more may help to get consistent structural connectome for ex vivo mouse brain studies.

### The effect of spatial resolution

Diffusion MRI at high spatial resolution offers an unprecedented power to assess the neural integrity and structural connectivity of the brain (Assaf, 2019; Calabrese et al., 2015; Huang et al., 2020; Vu et al., 2015; Wu and Zhang, 2016). Compared to conventional histology, it also affords a nondestructive way to explore the brain cyto- and myelo- architecture (Trampel et al., 2019; Wang et al., 2020; Yon et al., 2020). The advantages of high-resolution dMRI have been demonstrated in human brain studies (Chang et al., 2015; Setsompop et al., 2013; Vu et al., 2015), where the white matter tracts can be more accurately detected at submillimeter isotropic resolution (Hope et al., 2012). The HCP demonstrated that high-spatial resolution dMRI datasets (1.25 mm^3^ isotropic) provided great specificity and allowed reconstruction of certain tract features that were not observed by the lower spatial resolution (2 mm^3^ isotropic), even when the latter had higher angular resolution (Setsompop et al., 2013; Vu et al., 2015). Cortical anisotropy can also be observed in the HCP datasets (isotropic spatial resolution of 1.25 mm), but not in more conventional clinical datasets (isotropic spatial resolution of 2 mm) (Setsompop et al., 2013; Vu et al., 2015). A previous ex vivo macaque brain study showed that the prevalence of crossing fibers increased from 23% to 29% of all white matter voxels with an analysis of SNR-equivalent for various spatial resolution (300 to 800 µm), while this increase was much smaller than that found for increasing resolution (Schilling et al., 2017a). The SNR of the highest resolution MRI dataset (300 µm) was about 38 in the b0 image. The SNR of the b0 image of this study is more than 80 and the SNR of the DWI at b value of 8k s/mm^2^ is about 15, values much higher than the typical human DWIs (Chang et al., 2015; Wu and Miller, 2017). The MFR is dependent on the spatial resolution in mouse brain, which may help to explain the tractography and connectome results acquired at different spatial resolution. These findings may shed the light for developing new tractography algorithms as well as microstructural models with the consideration of the complex fiber structures in the brain (Schilling et al., 2017a; Schilling et al., 2018).

A few studies have been conducted to analyze the tractography variations at different b value and post-processing parameters, but the datasets were acquired at relatively low spatial resolution (Aydogan et al., 2018; Daianu et al., 2015). For instance, b value effect was accessed using a 5- shell HARDI (1000 – 12000 s/mm^2^) at 100 × 100 × 200 µm^3^ spatial resolution (Daianu et al., 2015). Another study at 200 µm isotropic resolution demonstrated that tractography results varied significantly with respect to the all the tracking parameters using multi-shell dMRI data from mouse brains and tracer injections from ABMA with extensive multidimensional tractography validation experiments (over 1 million) (Aydogan et al., 2018). The highest spatial resolution in our study is 25 µm^3^, which is at least 128 times higher than previous studies. Despite the high quality of the dMRI data presented here, its correspondence with neuronal tracer results is still not ideal. It is consistent with previous comparisons of neuronal tracer and diffusion based tractography (Aydogan et al., 2018; Calabrese et al., 2015). The viral tracers for ABMA follow individual axons and do not cross synapses. Tractography algorithms follow the bundles of axon and continue to propagate after synapses (Thomas et al., 2014). Conducting neuronal tracer injections and dMRI tractography on the same mice may further help to improve the correlations between these two techniques.

It has been reported that the neuroanatomy of the mouse brain can be better resolved at microscopic resolution, subtle fiber orientations and projections in hippocampus can also be better delineated using dMRI (Wang et al., 2020). However, the high spatial resolution dMRI requires extremely longer scan time (Jiang and Johnson, 2010). In a previous work, a dMRI dataset was acquired for about 10 days to achieve the isotropic resolution of 43 µm with 120 DWIs, which prevents any population studies in terms of both time and cost (Calabrese et al., 2015). To reduce the scan time, compressed sensing (CS) has been implemented in the 3D Stejskal-Tanner spin echo diffusion-weighted imaging sequence to reduce the scan time up to 8 times (Lustig et al., 2007; Wang et al., 2018a). Other fast acquisition techniques, such as parallel imaging, simultaneous MultiSlice (SMS) acquisition may further help to reduce the scan time (Barth et al., 2016; Deshmane et al., 2012; Glockner et al., 2005).

The adult human brain contains about 86 billion neurons, containing ∼ 62,732 neurons in 1 mm^3^ (Herculano-Houzel, 2009; Insel et al., 2013). The high spatial resolution dMRI in clinical scan can achieve 1mm^3^ or even submillimeter resolution, each imaging voxel still has thousands of neurons, which is too large to enable the resolution of axons (Buzsaki et al., 2013). The adult mouse brain is about 3000 times smaller than the human brain and it contains about 187,740 neurons in 1 mm^3^ (Herculano-Houzel, 2009; Johnson et al., 1993). The voxel size of mouse brain in this study is nearly 64,000 times smaller than the clinical scans, which results in about 3 neurons in a single imaging voxel. Compared to the brain volume and spatial resolution differences, the axon diameter difference between rodent brain and human brain is much smaller (Barazany et al., 2009; Lee et al., 2019; Liewald et al., 2014). The axon diameter is about 0.56 µm in control mice and about 0.62 µm in Rictor conditional knockout (CKO) mice (West et al., 2015, 2016). The axon diameter can be as small as 0.3 µm in the human brain, and as large as 20 µm, with a majority of thin fibers (∼ 84%) smaller than 2 µm (Liewald et al., 2014; Saliani et al., 2017). Considering the relatively small variation of the axon diameter between mouse and human brains, the improvement of spatial resolution in rodent MRI may dramatically reduce the partial volume effect and could be used to develop new tractography algorithms and validate novel diffusion biophysical models (Henriques et al., 2019; Huang et al., 2020; Palombo et al., 2020).

The mean cortex thickness of the mouse brain is 0.89 mm (with 36 voxels at 25um^3^), and the thickness in a somatosensory adult mouse cortical thickness is ∼ 1.7 mm (68 voxels at 25 um^3^). The human cerebral cortex is known to be a highly folded sheet of neurons with the thickness ranging from 1 to 4.5 mm (1 – 4.5 voxels at 1 mm^3^), with an overall average of approximately 2.5 mm (2.5 voxels at 1 mm^3^) (Fischl and Dale, 2000). The high spatial resolution in preclinical MRI may help to study the layer-specific cytoarchitecture variations of the cerebral cortex in different neurodegenerative diseases, which is still challenging in clinical studies due to the coarse resolution and partial volume effect (Assaf, 2019; Colgan et al., 2016; Vu et al., 2015; Wang et al., 2020; Wu and Zhang, 2016).

There are a few limitations to our study. An obvious limitation is the extremely long scan time for both high spatial resolution and angular resolution dMRI scans, which prevents to probing statistical differences between different acquisition protocols. The scan time can be largely reduced by reducing the number of shells, angular resolution, and/or spatial resolution to get comparable results to the fully sampled datasets (Wang et al., 2018a). Further reducing the scan time can be achieved by performing joint reconstruction of k-q space in future studies (Cheng et al., 2015). A more thorough comparison among different tractography algorithms and different software should be investigated in detail (Christiaens et al., 2015; Garyfallidis et al., 2018; Sarwar et al., 2019; Woolrich et al., 2009). Recently, quantitative comparison of the FODs have been proposed to provide more anatomically specific information about the microstructural properties of white matter populations, which is warrented in the future studies with sufficient sample numbers (Raffelt et al., 2017; Tournier et al., 2019; Wu et al., 2020).

Although ex vivo MRI can provide useful information on both the macroscopic (anatomical) and microscopic (microstructural) level, the results from ex vivo specimens may not accurately represent tissue microstructures under normal physiological conditions due to the significant microstructural changes associated with death and chemical fixation in post-mortem samples (Dyrby et al., 2011; Shatil et al., 2018). Diffusion scan with single-shot echo planar imaging (EPI) readout has been widely used for in vivo studies, EPI segmentation and partial Fourier encoding acceleration have been applied to reduce the echo time and distortion artifacts (Alomair et al., 2015). The macroscopic neuronal projection (100 µm isotropic) in the mouse brain hippocampus has been illustrated using 3D diffusion-weighted gradient- and spin-echo (DW-GRASE) based sequence (Aggarwal, et al, 2015; Wu and Zhang, 2016) with the b value of 2500 s/mm^2^, the effects of different acquisition parameters for in vivo mouse brain structural connectome still needs further investigation.

In conclusion, we systematically investigated the effects of different acquisition parameters to the final outcome of the structural connectome in ex vivo mouse brain. The structural connectome strongly depends on the b value, spatial resolution, and angular resolution. To achieve stable tractography and connectome results, more than 50 DWIs is essential for ex vivo mouse brain studies. A single-shell acquisition with b value of 3000 s/mm^2^ shows comparable results to the multi-shell acquisition. The dice coefficient decreases and both false positive rate and false negative rate gradually increase with coarser spatial resolution. The study may help to provide guidance for achieving stable tractography and structural connectivity results for ex vivo mouse brain studies.

## Acknowledgements

This work was supported by the NIH P41 EB015897, NIH 1S10OD010683-01, 1R01NS096720-01A1. The authors thank Lucy Upchurch and Brian Overshiner for significant technical support. The authors thank Dr. Yu-Chien Wu and Samuel Foust from Roberts Translational Imaging Facility and STARK Neuroscience Research Institute for resource support. The authors thank Tatiana Johnson for editorial comments on the manuscript.

## Compliance with ethical standards

### Ethical approval

All animal studies have been approved by the appropriate ethics committee: Duke University Institutional Animal Care and Use Committee.

### Informed consent

No human subject was used in this study.

## Supplemental Figures

**sFigure1:**
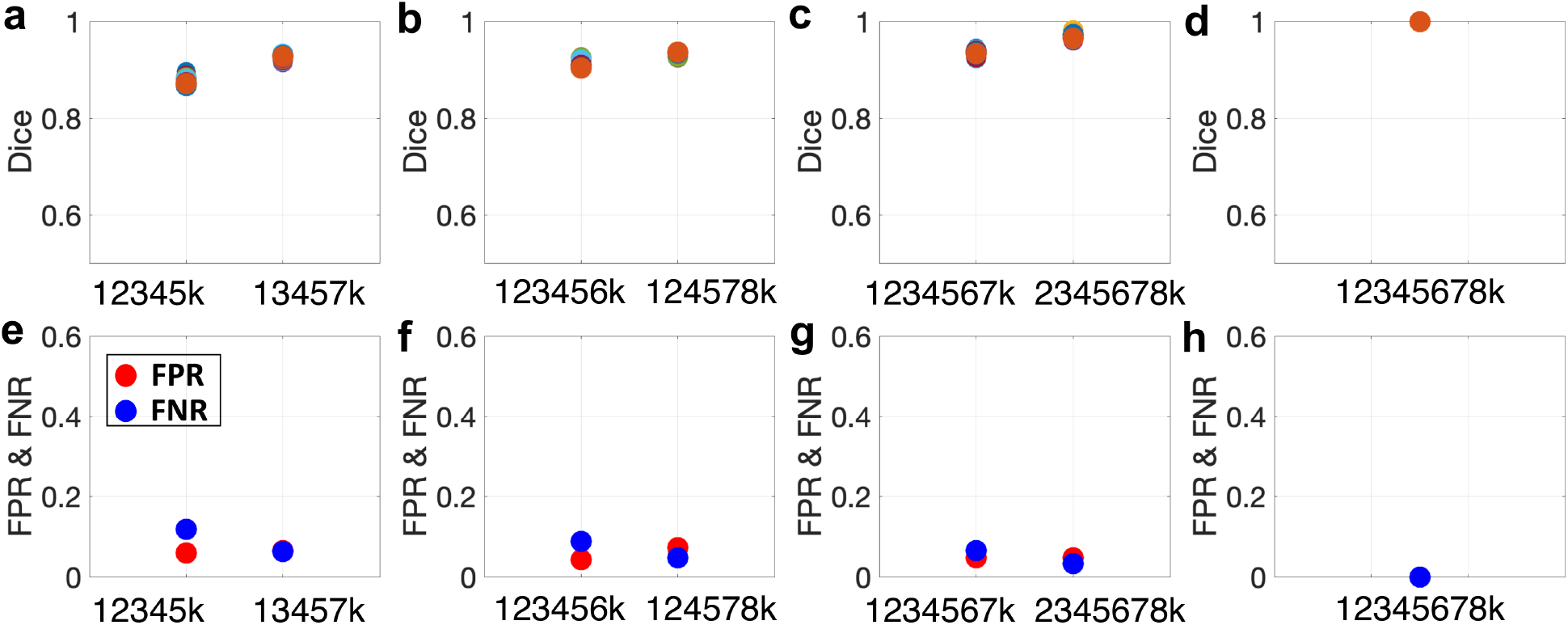
The tract density imaging (TDI) at different threshold (0.0001 – 0.2). The dice coefficient increases when more shells are added; the false positive rate (FPR) and false negative rate (FNR) decrease when more shells are added.

**sFigure2:**
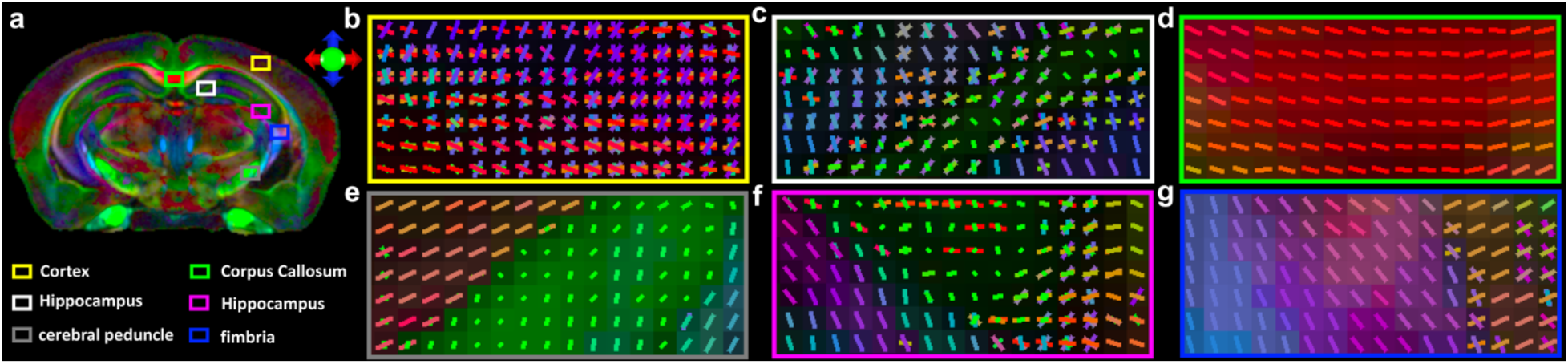
The color fractional anisotropy (FA) image (a) and complex fiber orientation distributions throughout the mouse brain, including cortex (b), hippocampus (c, f), corpus callosum (d), cerebral peduncle (e), and fimbria (g).

**sFigure3:**
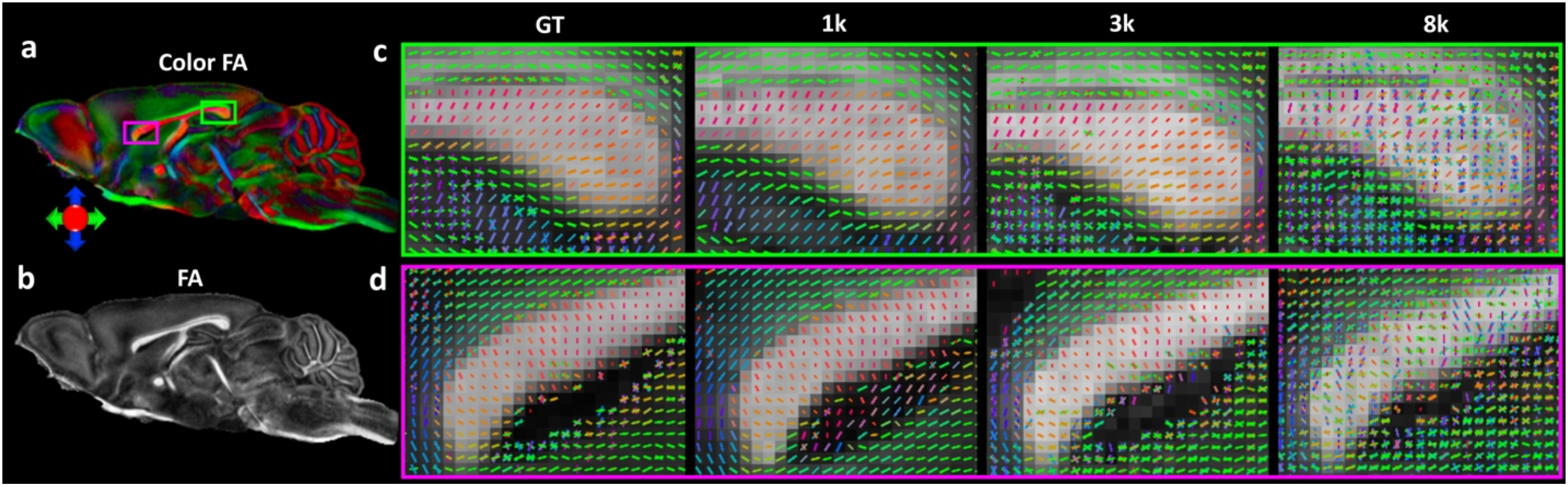
The fiber orientation distributions in the corpus callosum at different b values. Few crossing fibers are resolved at b value of 1000 s/mm^2^ and 3000 s/mm^2^, while crossing fibers are evident in most of the voxels at b value of 8000 s/mm^2^.

**sFigure 4:**
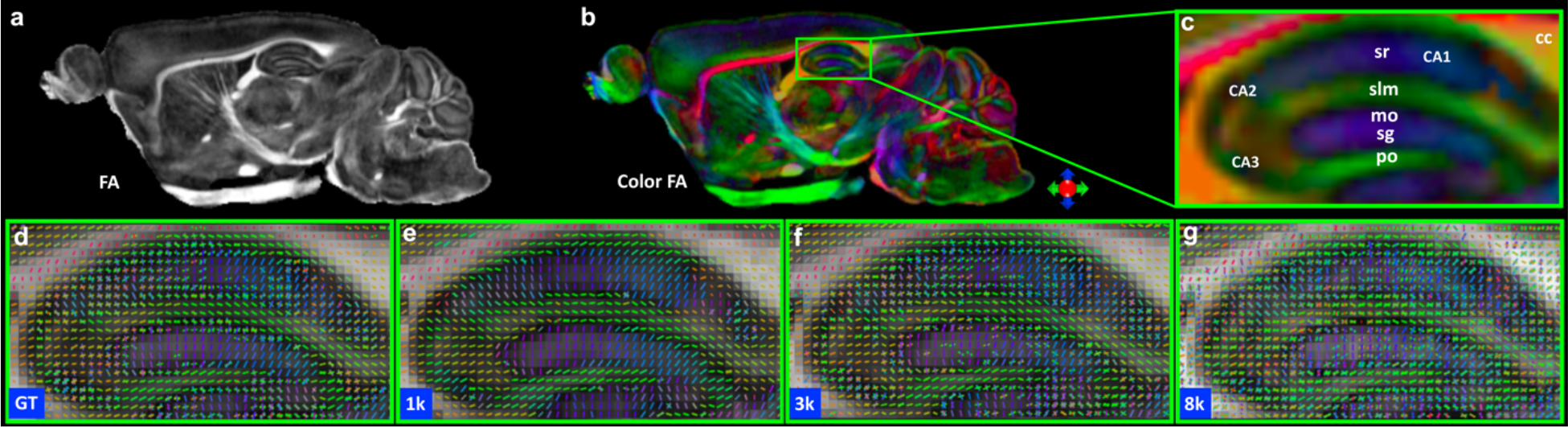
The b value dependent fiber orientation distributions of hippocampus. Different layers with distinct fiber orientations are demonstrated in hippocampus (b-c). Few crossing fibers are resolved at b value of 1000 s/mm^2^ (e), while crossing fibers are present in most the voxels in the hippocampus at b value of 8000 s/mm^2^ (g). The fiber orientation distributions at b value of 3000 s/mm^2^ (f) are visually comparable to the ground truth in the hippocampus (d). cc: corpus callosum; sr:stratum radiatum; slm: stratum lacunosum-moleculare; mo: molecular layer; sg: granule cell layer of the dentate gyrus; po: polymorphic layer of the dentate gyrus;

**sFigure 5:**
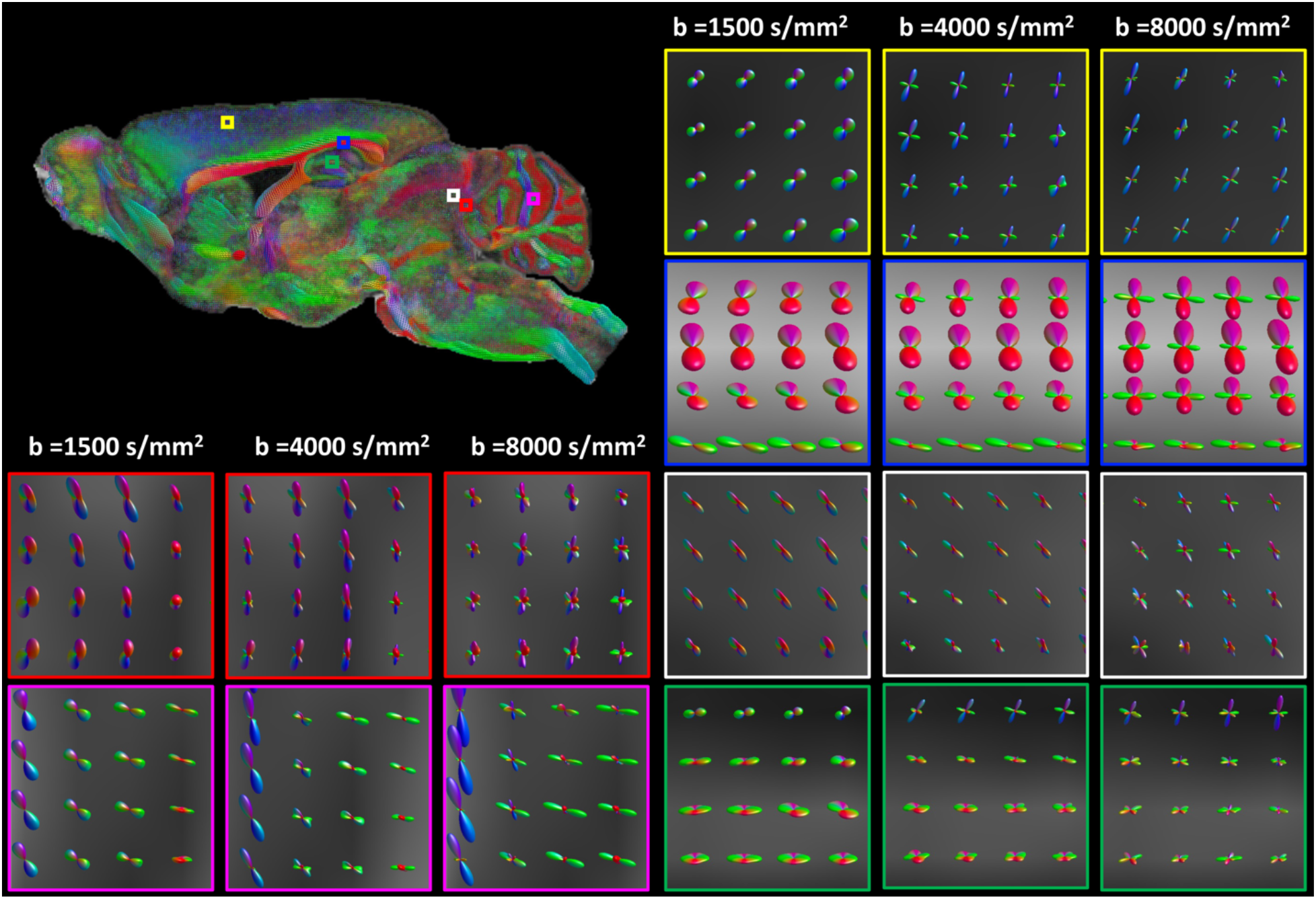
The b value dependent fiber orientation distributions of different regions of mouse brain using MRtrix3. Few crossing fibers are resolved at b value of 1500 s/mm^2^, while crossing fibers are dominated through the whole at b value of 8000 s/mm^2^.

**sFigure 6:**
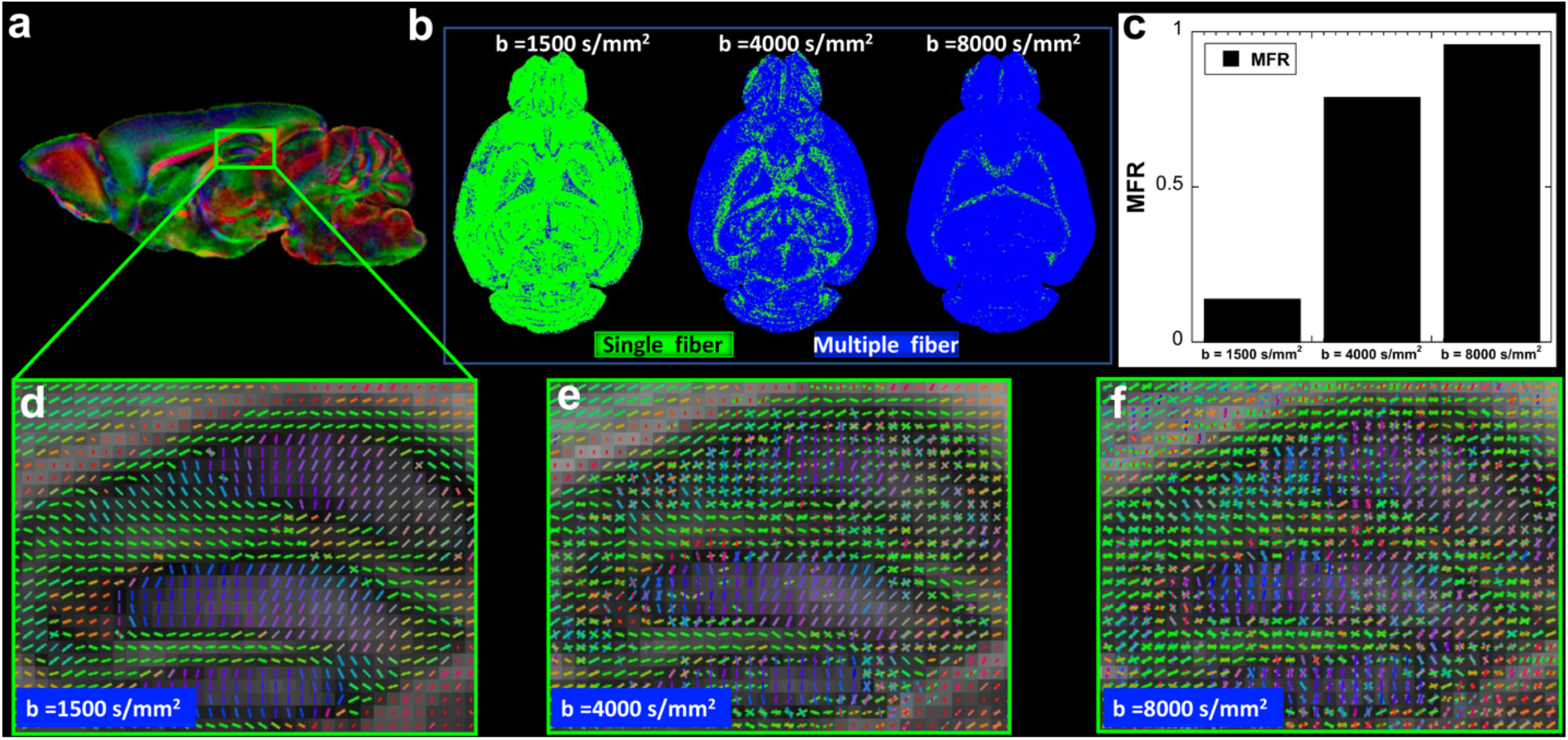
multiple fiber ratio (MFR) mapping at different b values. The MFR is extremely low at b value of 1500 s/mm^2^. The MFR gradually increases with b value and the MFR is close to 1 at b value of 8000 s/mm^2^ (c).

**sFigure 7:**
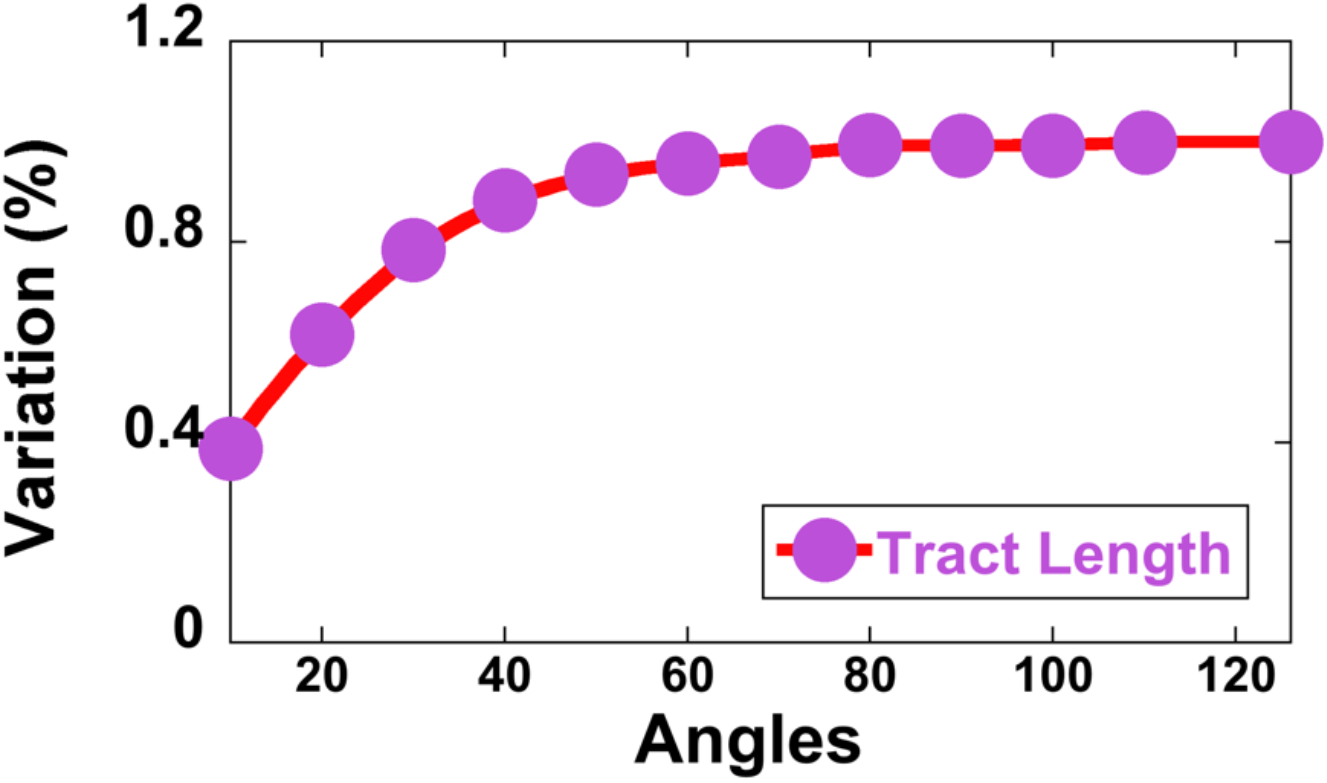
The tract length (TL) variations with angular resolution. The track length values are normalized to the TLvalue at 126 angular resolutions. The TL becomes stable when the angular resolution is higher than 50.

